# A translational swine model for Crohn’s disease

**DOI:** 10.1101/2022.07.08.499360

**Authors:** T Winogrodzki, A Metwaly, A Grodziecki, W Liang, B Klinger, K Flisikowski, T Flisikowska, K Fischer, K Steiger, D Haller, A Schnieke

**Author notes:** These authors contributed equally to this work (T Winogrodzki, A Metwaly, D Haller, A Schnieke). **Correspondence to:** Prof. Angelika Schnieke. Chair of Livestock Biotechnology. TU München. Liesel-Beckmann-Straße 1, 85354, Freising, Germany. **Writing assistance**: None. **DNA sequence data:** NCBI accession number GCF_000003025.6, Sscrofa 11.1. **Author Contributions:** T.W., K.Fli., D.H. and A.S. designed research; T.W., B.Kli., W.L., A.G., T.F., A.M. performed research; T.W., A.M., K.Fli., K. Fis. and A.S. analyzed data; T.W., K. Fli, T. Fli, K. Fis., K.S., D.H. and A.S. wrote the paper.

## Abstract

Crohn’s Disease (CD) is incurable, and represents a lifelong burden for patients and its incidence is increasing worldwide. A key contributing factor is a dysregulated immune response. Here we report the generation of genome edited pigs with a deletion of the transcript-destabilizing AU-rich element (ARE) and a constitutive decay element (CDE) in the *TNF* gene which recapitulate major characteristics of human CD, including ulcerative transmural ileocolitis, increased abundance of proinflammatory cytokines, evidence for impaired integrity of the intestinal epithelial cell barrier, immune cell infiltration, and dysbiotic microbial communities. This physiologically relevant CD model enables human-scale and long-term studies to assess diagnostic, nutritional or microbial interventions, filling the gap for translating findings into the clinic.

---

Crohn’s disease (CD) is one of the two main subtypes of inflammatory bowel diseases (IBD) affecting the gastrointestinal tract characterized by patchy transmural inflammation in the entire digestive tract, predominantly affecting the terminal ileum and proximal colon^1^. CD is a multifactorial disease driven by complex gene-environment interactions following patterns of industrialization^2^. Genome-wide association studies identified a variety of genetic risk alleles pointing towards a disruption of microbe-host interactions^3^. Underlying molecular mechanisms of IBD and specific microbial and metabolic signatures have been elucidated using spontaneous, chemically-induced and genetically engineered mouse models^4^, however CD-like inflammation with small intestinal disease manifestation rarely occurs in currently available IBD-related animal models (e.g. SAMP/YitFc, *Xiap-/-, Xbp1-/-*, *TNF^ΔARE^*).Hence, the need for additional animal models reflecting the human pathology more closely and enabling translational, e.g. dietary or microbial studies. Pigs are omnivores, gnotobiotic rearing is possible as well as long-term studies. The porcine immune system resembles that of humans more closely than the mouse^5^. Both porcine^6^ and human bacterial libraries^7^ are available to elucidate the effect of specific bacterial strains on IBD. Microbiome transfer from humans to pigs results in a gut microbiota closely resembling that of the human donor^8^. Novel diagnostic technologies can be assessed at human scale^9^.

To address the clinical need, we generated a porcine CD model based on the *TNF^ΔARE/+^* mouse^10^. Genome editing in *in vitro* produced embryos excised a 93 bp fragment in the 3’ untranslated region of the *TNF* gene containing an AU-rich element (ARE) and one of two constitutive decay elements (CDEs) (Figure 1A, S1A). ARE and CDEs interact with RNA-binding factors, their deletion reduces *TNF* mRNA decay, resulting in increasing TNF protein levels and intestinal inflammation. As homozygous deletion of ARE and both CDEs caused embryonic lethality in mice^11^, only one CDE was deleted from the porcine *TNF* gene. Two pregnancies resulted in the birth of ten piglets, of which two showed a correct, biallelic excision of the *ARE/CDE1* sequence (*TNF^ΔARE/ΔARE^*, pig# 2, 5), and five a monoallelic deletion (*TNF^ΔARE/+^*, pig# 1, 3, 6, 8, 9) (Figure S1A, B). Two of the latter piglets (#8, 9) showed mosaicism and/or InDel mutations on the second allele, which was eliminated from the line after breeding with wild-type animals. Founder animals (# 6, 8, 9), used for breeding, were analyzed for the five most likely off-targets, none were detected. Breeding resulted in 11 *TNF^ΔARE/+^*F1 and ten F2 offspring (7 *TNF^ΔARE/+^*, 3 *TNF^ΔARE/ΔARE^*).

**Figure 1.**
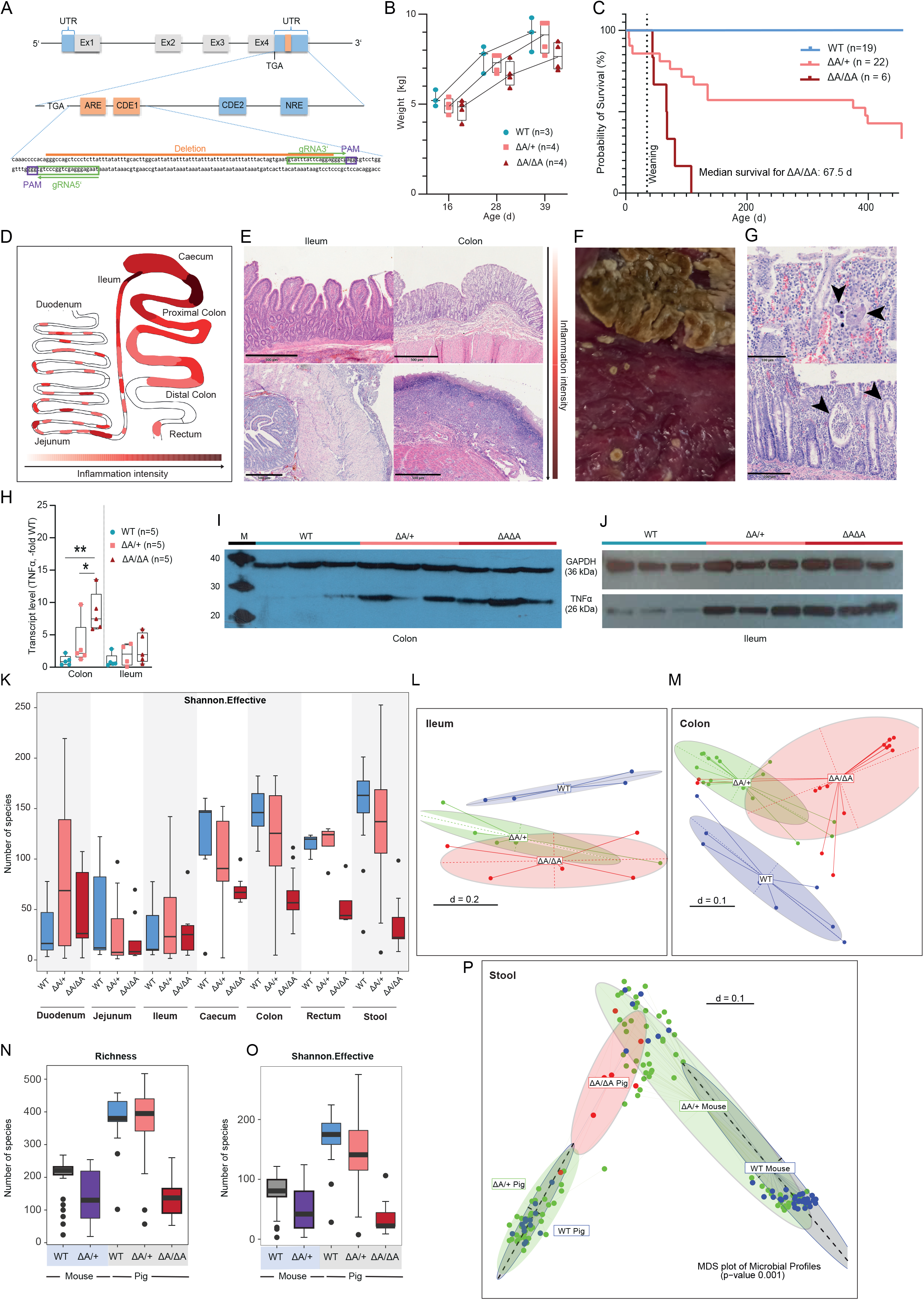
(A) Gene Targeting strategy for the deletion of the ARE and CDE1 elements. Shown within the 3’-untranslated region (UTR) are miRNA- and RNA-binding protein sequence motifs, known to play a role in mRNA decay: ARE, CDE and NRE. gRNA binding sites and their PAM sequences are highlighted. (B) Comparison of weight gains of pigs from the same litter at 16 to 39 days of age. (C) Kaplan-Meier curve of the time at which a tolerable health score was passed over and required euthanasia of the animals. The median survival time for TNF^ΔARE/ΔARE^ pigs is given. (D) Schematic representation of the different sites of inflammation. A darker red color indicates a more intense inflammatory process. (E) Representative images of H&E stained colonic and ileal gut sections. Shown are mildly inflamed (top) and highly inflamed (bottom) tissue sections with crust formation (colon) and serositis (ileum). Bars indicate 500 μm. (F) Diphteritic membranes in the caecum of a TNF^ΔARE/ΔARE^ pig. (G) Histopathological findings of invasive Balantidium coli (top) and crypt abscesses (bottom). Bars indicate 100 μm (top) and 200 μm (bottom), respectively. (H) Comparison of TNF-transcript levels of colonic and ileal mucosal biopsy samples. Transcript levels are presented as multiples of the wild-type average value. (I, J) Western blot analysis of TNF and GAPDH of colonic and ileal samples from wild-type, 3 TNF^ΔARE/+^ and 3 TNF^ΔARE/ΔARE^pigs. MagicMark Western Protein standard (Invitrogen) and BlueRay Prestained Protein Marker (Nippon) were used as standard ladders (M). (K) Alpha-diversity of the luminal and mucosa-associated microbiota. Richness and Shannon effective number of species. (L, M) MDS plot of microbial profiles of ileal and colonic samples stratified by genotype, respectively. (N, O) Alpha-diversity of luminal microbiota from wild-type and TNF^ΔARE^pigs and mice shown as richness and Shannon effective number of species. (*P*) MDS plot of microbial profiles fecal microbiota stratified by species and genotype.

After weaning ~50% of *TNF^ΔARE/+^* animals showed reduced weight gain and reduction in stool consistency. This phenotype was more pronounced in *TNF^ΔARE/ΔARE^* pigs (Figure 1B, C). Affected heterozygous and homozygous pigs were euthanized when severe symptoms (chronic diarrhea, severe weight loss, apathy) manifested (Figure 1C). Macroscopically, animals showed intestinal edema and hemorrhage throughout the colon, segmental in the small intestine, frequently including the caecum (Figure 1D and S1C, D). As with mice, inflammation intensity varied between pigs of the same genotype (Figure 1E). In *TNF^ΔARE/ΔARE^* pigs, the strongest observed alteration was ulcerative enteritis, sometimes covered with diphteritic membranes in the caecum (Figure 1F). Microscopically, 67% (4/6) of pathologically evaluated *TNF^ΔARE/+^* animals showed ileitis and/or colitis with mixed infiltrations of the lamina propria often extending into the tela submucosa accompanied by (fibrino-)suppurative serositis (Figure 1E). In rare cases, extensive crypt abscesses (Figure 1G) and frequent herniation of crypts throughout the lamina muscularis mucosae or ulceration of the lamina propria mucosae with mixed infiltration and development of a fibroangioblastic granulation tissue were observed. Overall *TNF^ΔARE/ΔARE^* pigs showed more severe intestinal alterations, up to severe ulcerative to necrotizing inflammation with intraluminal accumulation of cellular debris. In one case, also invasion of *Balantidium coli* into the mucosa was observed (Figure 1G). Immunohistochemistry confirmed increased cell proliferation in affected areas (Ki67), increased leukocyte infiltrations (CD3^+^, IBA1^+^) and decreased number of mucus-secreting goblet cells, especially in the ileocolonic region (Figure S1E,F). Complete blood count of these pigs showed increased monocyte count without significant change in total white blood cells, basophilia, increased urea-creatinine ratio and hypoalbuminemia indicative of gastrointestinal bleeding^12,13^, elevated serum Cu/Zn ratio associated with malnutrition and inflammation, which has been proposed as a potential bio-marker for IBD patients^14^ (Figure S1G, H).

To confirm that altered expression of TNF and its downstream targets (e.g. IL6; IL8), was causing the pathophenotype, mRNA half-life, mRNA expression and protein levels were assessed. For the determination of TNF-mRNA half-life, macrophages were cultured, stimulated with LPS and treated with actinomycin D. The resulting calculated half-life was ~19 min for wild-type, ~58 min for *TNF^ΔARE/+^* and ~708 min *TNF^ΔARE/ΔARE^*samples (Figure S1I, J). Compared to wild-type increased *TNF* mRNA and protein expression was detected in the colon and ileum of *TNF^ΔARE^* animals (Figure 1H-J). This was accompanied by an upregulation of the key pro-inflammatory cytokine IL6 and the neutrophil chemoattractant IL8 (Figure S1K).

Microbiota composition was analyzed by 16S rRNA gene amplicon sequencing of 61 stool and 166 mucosal samples from defined positions along the intestinal tract (7 wild-type, 16 *TNF^ΔARE/+^* and *TNF^ΔARE/ΔARE^* pigs) and cross-species comparison with the mouse^15^ was performed. Alpha-Diversity was reduced in large intestine and jejunum of TNF^ΔARE^ pigs, while it was enriched in duodenum and ileum (Figure 1K). Beta-diversity analysis showed separation of microbial profiles in ileal and colonic mucosa of wild-type, *TNF^ΔARE/+^* or *TNF^ΔARE/ΔARE^* pigs, which was weakly reflected in stool. Members of the phyla Proteobacteria or Fusobacteriota and Campylobacteiota were increased in stool, ileal or colonic-associated microbiota of TNF^ΔARE^ pigs (Figure 1L, M and S2A-F). These as well as the identified set of bacterial genera (Figure S2G-I), including members of the *Megasphaera*, were previously linked to human IBD^16^,^17^. Overall outbred pigs showed significantly higher bacterial community richness and diversity compared to inbred mice, but *TNF^ΔARE/ΔARE^* pigs showed considerably reduced number of species (Figure 1N, O). The fecal microbiota from a subset of *TNF^ΔARE/ΔARE^* pigs clustered with *TNF^ΔARE/+^* mice, suggesting a potential microbiota clustering based on the severity of inflammation (Figure 1P).

In summary: The first large animal model for IBD recapitulates major characteristics of human Crohn’s disease, including ulcerative transmural ileocolitis, increased abundance of proinflammatory cytokines, evidence for impaired integrity of the intestinal epithelial cell barrier, immune cell infiltration, and dysbiotic microbial communities. This physiologically relevant Crohn’s disease model enables human-scale and long-term studies to assess diagnostic, nutritional or microbial interventions, filling the gap for translating findings into the clinic.

## Abbreviations used in this paper

IBD: Inflammatory bowel disease
CD: Crohn’s disease
TNF: Tumor necrosis factor alpha
ARE: AU-rich element
CDE: Constitutive decay element
Ki67: Marker of proliferation Ki67 (Kiel-67)
CD3: Cluster of differentiation 3
IBA1: Allograft inflammatory factor 1
IL6: Interleukin-6
IL8: Interleukin-8

## Acknowledgments

The authors would like to thank Lea Radomsky, Sara Robin and Monika Heilmeier for their technical support and Viola and Steffen Löbnitz for animal husbandry.

---

**Supplementary Figure 1.**
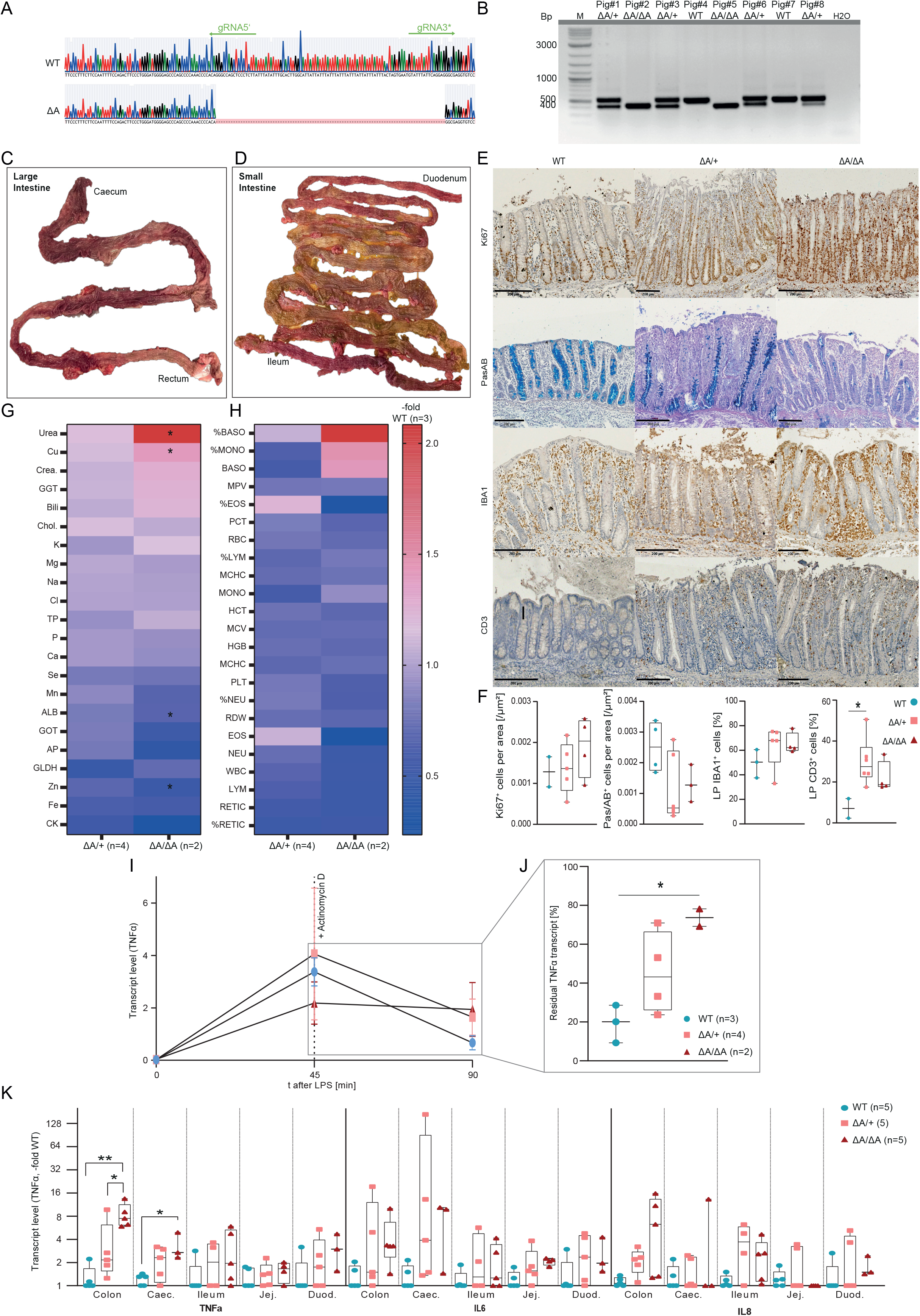
(A) Genotyping PCR of one founding litter (Pigs# 1-8) revealed the presence of a minor amplification product resulting from the 93 bp deletion. The presence of only a single band displays a wild-type (~ 550 bp) or TNF^ΔARE/ΔARE^ (~450 bp) genotype. The presence of two bands displays a TNF^ΔARE/+^ genotype. 2-log marker (M) was used as DNA ladder. (B) Sanger sequencing revealed a 93 bp deletion between the two gRNA binding sites. All TNF^ΔARE^ animals shared the same mutant allele sequence. (C, D) Representative finding of inflamed large and small bowel. While the macroscopically visible tissue coloration was continuous in the large intestine, it was segmental in the small intestine. (E, F) Inflammation-induced epithelial changes and aberrant mucosal architecture were detected in inflamed mutant pigs. Ki67^+^-, CD3^+^- and IBA1^+^-cells were enriched, while Pas/AB^+^-cells were depleted in intestinal crypts and lamina propria of TNF^ΔARE^ pigs compared to wild-type littermates. Shown are representative images and statistical analyses of colon sections. Scale bars indicate 200 nm. (G, H) Heatmap of the complete blood count of TNF^ΔARE/+^ and TNF^ΔARE/ΔARE^ pigs compared to wild-type littermates. Significantly differing levels are indicated. (GOT: glutamate-oxalacetate-transaminase; GLDH: glutamate-dehydrogenase; GGT: y-Glutamyl-Transferase; Billi: Bilirubin; AP: alkaline phosphatase; Chol.: cholesterol; TP: total protein; ALB: albumin; RBC: red blood cells; HCT: haematocrit; HGB: haemoglobin; MCV: mean corpuscular volume; MCHC: Mean corpuscular haemoglobin concentration; RDW: red cell distribution; RETIC: reticulocyte; WBC: white blood cell; NEU: neutrophils; LYM: lymphocytes; MONO: monocytes; EOS: eosinophils; BASO: basophils; PLT: platelets; MPV: mean platelet volume; PCT: procalcitonin). (I, J) Residual TNF transcript levels in in vitro macrophage culture after 45 minutes of LPS stimulation followed by 45 minutes of transcriptional inhibition by actinomycin D compared to 45 minutes of LPS stimulation only. (K) Comparison of TNF, IL-6 and IL-8 transcript levels in intestinal samples.

**Supplementary Figure 2.**
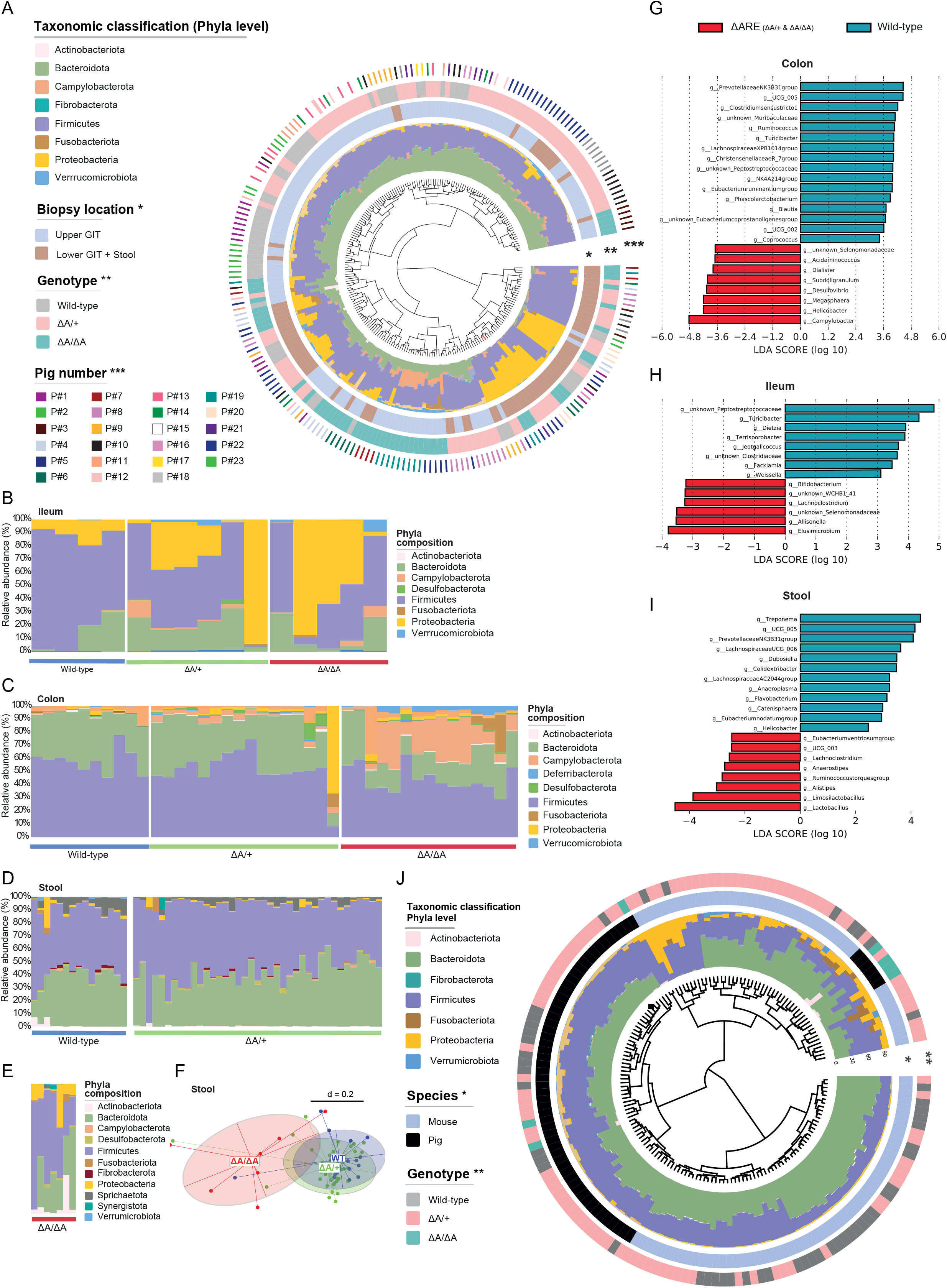
(A) Phylogenetic tree showing the similarities between microbiota profiles based on generalized UniFrac distances in luminal (n= 61) and mucosa-associated microbiota (n=166) derived from 23 pigs (7 wild-type, and 16 TNF^ΔARE^). Individual taxonomic composition at the phylum level is shown as stacked bar plots around the phylogram. Innermost ring shows stratification based on sample type, upper GIT samples (blue) and lower GIT and stool samples (brown) and denoted by (*), second ring shows stratification based on genotype, wild-type (grey); TNF^ΔARE/+^ (pink) and TNF^ΔARE/ΔARE^ (green) and denoted by (**). Bars in the outer ring of the figure indicate samples derived from each pig and denoted by (***). (B, C, D, E) Taxonomic composition at the phylum level in ileal, colonic and stool samples stratified by genotype. (F) MDS plot of microbial profiles of stool samples stratified by genotype. (G, H, I) Linear discriminant analysis effect size (LEfSe) indicating differentially enriched bacterial groups at the genus level of ileal, colonic and stool microbiota based on genotype. Red: TNF^ΔARE^ and Blue: wild-type. (J) Phylogenetic tree showing the similarities between microbiota profiles based on generalized UniFrac distances in luminal microbiota derived from wild-type and TNF^ΔARE^ pigs and mice. Individual taxonomic composition at the phylum level is shown as stacked bar plots around the phylogram. Innermost ring shows stratification based on species; mouse or pig and the outer ring shows stratification based on genotype, wild-type (grey); TNF^ΔARE/+^ (pink) and TNF^ΔARE/ΔARE^ (green).

## Methods

### Ethics statement

Animal experiments were approved by the Committee on Animal Health and Care of the local government body of the state of Upper Bavaria (ROB 55.2-2532.Vet_02-18-56, 55.2-1-54-2531-99-13) and performed according to the German Animal Welfare Act and European Union Normative for Care and Use of Experimental Animals.

### Generation of TNF^ΔARE^-pigs

For CRISPR/Cas9 mediated excision of 93 bp fragment containing the ARE and CDE1, two sgRNAs) were designed using CRISPOR^1^. Both U6-gRNA-scaffold sequences were cloned into a px330-U6-Chimeric_BB-CBh-hSpCas9 plasmid^2^ (Addgene #42230) (pX330-2gRNAs). The pX330-2gRNAs was microinjected into *in vitro* fertilized porcine oocytes followed by laparoscopic embryo transfer, as described^3,4^. Ear clip-gDNA was isolated using GenElute Mammalian Genomic DNA Kit (Sigma). The deletion was screened by PCR using GoTaq polymerase (Promega) and flanking primers ~190 bp distal to the gRNA target sites (Fwd: 5’-GGGTTTGGATTCCTGGATGC-3’, Rev: 5’-GCGGTTACAGACACAACTCC-3’). Thermal cycling parameters: 95°C, 2 min.; (35x) 95°C, 45 sec., 60°C, 45 sec., 72 °C, 30 sec.; 72 °C, 5 min. Amplicons were verified by Sanger sequencing. gRNA-off-target sites were predicted using CRISPOR. Five highest scoring potential off-targets were analyzed by PCR and Sanger sequencing.

### Macrophage culture and RNA half-life analysis

EDTA-blood was collected from anesthetized pigs. Peripheral blood mononuclear cells (PBMCs) were isolated by Ficoll-density gradient centrifugation. PBMCs were cultured in 10 ml RPMI 1640, 10 % FCS, 1 % GlutaMax (Sigma), 1 % Pen-Strep/Amphotericin B and 10^4^ U/ml recombinant poGM-CSF (Biotechne) for seven days. Macrophages were divided into groups: (1) Control, time points 0, 45, and 90 min; supplemented with: (2) 0.1 μg/ml LPS (InvivoGen) for 45 and 90 min; (3) LPS and 150 μg/ml Polymyxin B (Invivogen) for 45 and 90 min; (4) LPS for 90 min and 10 μg/ml Actinomycin D (Calbiochem) after 45 min. Macrophages were pelleted and frozen at −80 °C until processing.

### RNA isolation and quantification

RNA was isolated from gut mucosal biopsies using Monarch Total RNA Miniprep Kit (NEB) according to the manufacturers’ protocols. cDNA was generated using LunaScript RT Master Mix Kit (NEB) according to the manufacturer’s protocol. Real-Time qPCR was performed using the Fast SYBR Green Master Mix (Sigma) in a QuantStudio 5 Real-Time-PCR-Cycler (Thermofisher Scientific) (TNF Fwd: 5’-GGGCTTATCTGAGGTTTGAG-3’, Rev: 5’-TTCTGCCTACTGCACTTCGA-3’; IL6 Fwd: 5’-TCTGCAATGAGAAAGGAGATGTG-3’, Rev: 5’-AGGTTCAGGTTGTTTTCTGCC-3’; IL8 Fwd: 5’-CTGTGAGGCTGCAGTTCTG-3’, Rev: 5’-GTGATTGAGAGTGGACCCCA-3’). For transcript normalization, housekeeping genes (GAPDH Fwd: 5’-TTCCACGGCACAGTCAAGGC-3’, Rev: 5’-GCGGTTACAGACACAACTCC-3’); ß-Actin Fwd: 5’-TCCCTGGAGAAGAGCTACGA-3’, Rev: 5’-GCAGGTCAGGTCCACAAC-3’; RPS28 Fwd: 5’-GTTACCAAGGTTCTGGGCAG-3’, Rev: 5’-CAGATATCCAGGACCCAGCC-3’) were selected based on NormFinder^5^ and BestKeeper^6^. One-Way ANOVA was performed, followed by Tukey test for statistical evaluation using GraphPad Prism 8.

### TNF protein quantification

Ileal and colonic proteins were obtained by tissue homogenization in NP-40 Buffer with 1x cOmplete Mini Protease Inhibitor Cocktail (Roche). Protein concentrations were determined using Bradford Assay. Protein lysates were separated by semi-dry western blot (anti-TNF, 1:1000, Invitrogen (14-7321-85); anti-GAPDH, 1:3000, Sigma (G8795)). Blots were developed using Pierce ECL Plus Western Blotting Substrate (Thermo Scientific).

### Gross evaluation, histological examination and immunohistochemistry

Gross findings during necropsy were documented and images were reviewed by a board-certified veterinary pathologist (KS). Representative specimens for histology were collected from rectum, proximal colon, caecum, distal ileum, proximal jejunum, and proximal duodenum, fixed in 10% neutral-buffered formalin and embedded in paraffin (FFPE). Pathological evaluation was performed on 2 μm H&E stained tissue sections. Histopathological findings were described, morphological diagnoses were compiled on the slide level and the histophenotype findings were interpreted with respect to genotypes. Immunohistochemical stainings were performed on 3.5 μm FFPE tissue sections as described previously^7^ (anti-Ki67, 1:300, Invitrogen (MA5-14520); anti-CD3, 1:100, Southern Biotech (4511-01); anti-MPO, 1:300, Dianova (DLN-012930); anti-IBA1, 1:2000, Wako FujiFilm (019-19741)). Periodic acid-Schiff-Alcian blue (Pas/AB) staining was performed as described^7^. Quantification of lamina propria CD3^+^, IBA1^+^, and Ki67^+^ cells was performed by randomly selected areas and the ratio of positive/negative cells. Pas/AB^+^ and crypt Ki67^+^ cells were quantified counting positive cells/crypt area. Statistical evaluation was performed as described above.

### TNFα^ΔARE^ pig samples for microbiota profiling

Littermate pigs were co-housed at the Technical University Munich. Antidiarrheal medication was prohibited. Animals received the same diet (HEMO U 134 pellets (LikraWest)) and water ad libitum. Feces were collected monthly by digital sampling. Bristol stool score was determined during sampling. Gut luminal and mucosal tissue samples were collected during necropsy as described above.

### TNF^ΔARE^ mouse samples for microbiota profiling

TNF^δARE^ mice and wild-type control littermates were imported from Case Western Reserve University, Cleveland, and housed under germ-free (GF) conditions. 8 weeks old GF mice were colonized with specific-pathogen-free (SPF)-derived wild-type cecal microbiota and maintained under SPF housing conditions. Fecal samples were collected over time (8, 10, 12, and 18 weeks) and frozen at −20°C for downstream analysis.

### High-Throughput 16S rRNA Gene Amplicon Sequencing

Metagenomic DNA was extracted as described^8^. Briefly, cells were mechanically lysed in DNA stabilization buffer and phenol/chloroform/isoamyl alcohol (25:24:1, by vol.) using a bead-beater. After heat treatment and centrifugation, supernatants were treated with RNase. DNA was purified with NucleoSpin gDNA Clean-up Kit (Macherey-Nagel), following the manufacturer’s instructions. Amplification with primers 341F-ovh and 785r-ovh^9,10^ and sequencing of the V3/V4 region of 16s rRNA genes was performed as described^11^. 227 samples were sequenced in paired-end modus (PE275) using a MiSeq system (Illumina) according to the manufacturer’s instructions.

### Amplicon sequence analysis

Raw 16s rRNA amplicon reads were pre-processed using the Integrated Microbial Next Generation Sequencing pipeline^12^. Five nucleotides on the 5’ end and 3’ end were trimmed for the R1 and R2 read (trim score 5) and an expected error rate of 1. Detected chimeric sequences were removed using UCHIME^13^. Sequences with relative abundance <0.25% and <300 and >600 nucleotides were excluded from analysis. A zero-radius OTUs (zOTUs) table was constructed considering all reads before quality filtering^14–16^. Downstream analysis was performed using Rhea^14^. Taxonomy assignment was done using RDP classifier version 2.11 and confirmed using SILVA database^15^. For phylogenetic analyses, maximum-likelihood trees were generated by FastTree based on MUSCLE alignments in MegaX^16^. Alpha-Diversity analysis was computed using community richness and Shannon effective number of species. Beta-diversity analysis was performed using the generalized UniFrac distances. Permutational multivariate analysis of variance (PERMANOVA) was performed for statistical evaluation of beta-diversity.

